# Honeybees acquire a spatial memory by learning landmarks and visual patterns in parallel

**DOI:** 10.1101/2025.09.25.678702

**Authors:** Nicolas Scheuring, Keram Pfeiffer, M. Jerome Beetz

**Author notes:** Correspondence (M.J.B.).

## Abstract

Spatial memory is vital for central-place foragers that navigate the same habitat on a daily basis. Computational models suggest that insects remember panoramic snapshots to return to goal locations. To what extent these snapshots represent retinotopic images or whether salient landmarks are stored isolated from the panorama remains unclear. With a newly designed spatial-memory task, we show that honeybees employ two different strategies for learning visual cues. In essence, they learn visual patterns as panoramic snapshots and three-dimensional objects as single entities. Freely walking honeybees were challenged to locate a rewarded feeder in the presence of visual patterns or objects. Selectively perturbing the panorama degraded the orientation behaviour only when visual patterns were present but not when objects were available. Presenting visual patterns and objects simultaneously and setting these cues in conflict revealed that honeybees primarily rely on a landmark-based navigation strategy. Making the spatial information of the objects ambiguous did not induce disorientation because the honeybees then relied on the visual patterns for finding the feeder. Altogether, our results suggest two independent visual memories operating in parallel, one dedicated to local landmarks and a second one for visual patterns that could be interpreted as distal cues. Notably, in rats, the spatial tuning of some hippocampal cells has been shown to be either anchored to local or distal cues. Our results provide the first hints that a distinction between local and distal cues may also be present in the insect brain.

## Introduction

For visual navigation, insects often rely on celestial and terrestrial cues (Collett and Graham, 2004, Collett et al., 2013, Freas and Cheng, 2022). While celestial cues such as the sun are used for a compass sense (Grob et al., 2017, Dacke and el Jundi, 2018, Nguyen et al., 2021, Beetz et al., 2022, Beetz et al., 2023), terrestrial cues are site specific and can be used to encode places (Tinbergen and Kruyt, 1937, Narendra and Ramirez-Esquivel, 2017, Konnerth et al., 2023). Place coding is of particular importance for central-place foragers such as bees and ants that begin their foraging trip at a fixed location, i.e., their nest. Remembering the locations of flower patches is key for efficient foraging trips (Woodgate et al., 2017, Buatois and Lihoreau, 2016, Lacombrade et al., 2025, Moura et al., 2023). Early studies revealed that insects are guided by salient landmarks such as trees or rock formations (von Frisch and Lindauer, 1954, Wehner and Raber, 1979, Tinbergen and Kruyt, 1937). In addition to landmarks, panoramic scenes also guide insects (Cartwright and Collett, 1983, Judd and Collett, 1998, Fukushi, 2001, Graham and Cheng, 2009, Buehlmann et al., 2016, Towne et al., 2017). Changing the visual panorama degrades the orientation capabilities of ants (Wystrach et al., 2011, Buehlmann et al., 2016, Narendra and Ramirez-Esquivel, 2017). To increase the robustness of navigation, it has been discussed that insects store multiple panoramic snapshots along their foraging route (Cartwright and Collett, 1983, Collett et al., 2013). For example, while exploring, ants occasionally rotate and often stop rotating when facing their goal, i.e., their nest or a food spot (Nicholson et al., 1999, Fleischmann et al., 2016). These visual gaze fixations have been interpreted as time points when panoramic snapshots are stored (Judd and Collett, 1998, Zeil et al., 2003, Collett et al., 2013, Fleischmann et al., 2016, Grob et al., 2017). Because of the link between goal direction and the stored snapshot, insects head to the goal in subsequent trips as soon as the current view matches the stored snapshots (image-matching) (Cartwright and Collett, 1983, Harris et al., 2007, Collett, 2010, Zeil, 2012, Collett et al., 2013, Durier et al., 2003).

Despite extensive research, it is unclear how the insect brain stores a visual scene, i.e., whether it is sufficient to remember salient objects (landmark-based navigation). Another hypothesis is to store a retinotopic image that could be used for a pixel-wise comparison with the current view (Cartwright and Collett, 1983, Franz and Mallot, 2000, Collett et al., 2013, Webb and Wystrach, 2016, Webb, 2019). While behavioural results indicate that ants store a panoramic image in which salient objects are part of the panorama (Wystrach et al., 2011), another study has shown that bumblebees store local objects as landmarks isolated from visual patterns in the background (Doussot et al., 2020). These results in bumblebees were explained with a computational model assuming the storage of multiple snapshots, for example, one that contains exclusively the visual patterns, and a second one purely based on local objects. Here, we designed a behavioural experiment that tested whether honeybees store three-dimensional objects as single entities – which we refer to as local landmarks from here on – or as being anchored to the panorama. In contrast to visual patterns that were learned as a panoramic snapshot, our results, surprisingly, show that honeybees learned the local objects as landmarks. Moreover, honeybees learned the local landmarks and the visual patterns in parallel and could dynamically choose which visual memory to use for navigation.

## Results

### Visual place learning in walking honeybees

To study visual place learning in honeybees, we designed a cylindrical arena (40 cm in diameter, Figure 1A). On the arena’s floor, two visually identical feeders were placed, but only one feeder was rewarded. Honeybees were trained to find the rewarded feeder with the aid of visual patterns presented at the arena’s inner wall. The visual patterns consisted of a ‘T’ and a ‘Ʇ’ (Figure 1A). Notably, these patterns were 90° offset to the feeders to avoid beaconing for reaching the rewarded (correct) feeder. To begin a trial, a wing-clipped honeybee was released at the arena’s centre (Video S1) and she was allowed to forage freely until probing one of the two feeders. Afterwards, the honeybee was caught with an opaque cup and returned to the arena’s centre. To avoid an orientation based on possible scent marks from previous trials, we moved the feeders and the visual patterns before releasing the honeybee to begin the next trial.

**Figure 1.**
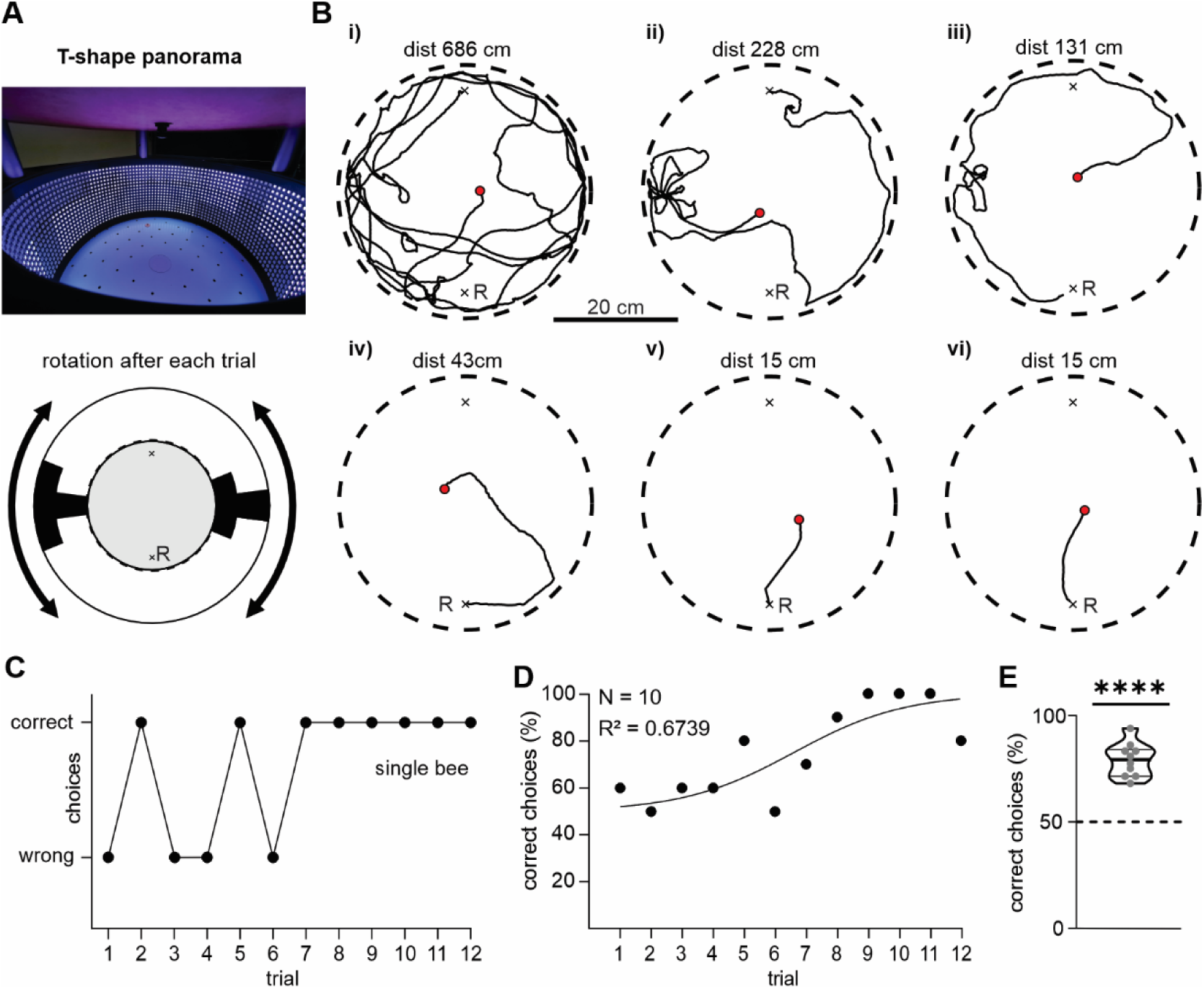
Honeybees can use visual patterns for place learning. (A) Photo and schematic top view of the arena. (B) Representative trajectories of one honeybee over six trials (i-vi). Trials begin at the red dot and end when the animal probed a feeder (x = unrewarded; R = rewarded). (C) Choices of one honeybee over twelve trials (correct = rewarded feeder). (D) Averaged data from ten animals fitted with a logistic model (N = 10; R^2^ = 0.6739). (E) Data points from (D) summarized in a violin plot. Dashed horizonal line indicates chance level (t = 14.15, p < 10^-5^, N = 10; one sample t-test).

Across trials, honeybees less often visited the unrewarded (wrong) feeder and travelled shorter distances until probing a feeder (Figure 1B, 1C; Video S2). The increase of correct choices for ten animals resembled a logistic model (Figure 1D; N = 10; R^2^ = 0.6739). Altogether, the honeybees chose the correct feeder above chance level (Figure 1E; t = 14.15, p < 10^-5^, N = 10; one sample t-test).

### Honeybees demonstrate image matching

After showing that honeybees learned the spatial relation between the visual patterns and the correct feeder, we next asked which part of the visual patterns the animals had learned. One possible strategy is to learn only one pattern. For example, honeybees faced the correct feeder when keeping the ‘T’ in the right visual field. Alternatively, honeybees could have learned the entire panoramic scene. Thus, after every three training trials, we added a vertical stripe (Figure 2A; *test*). Importantly, the stripe exclusively perturbed the panoramic view but not the ‘T-shapes’. If the honeybees had learned a panoramic snapshot, then they should be disoriented during test trials. Indeed, compared to training, bees showed longer and more tortuous walking trajectories during test trials (Figure 2A). In addition, the number of incorrect choices increased during test trials (Figure 2B, 2C; W = 36, p = 0,0078; N = 9; Wilcoxon matched pairs signed rank test). The rate of correct choices was above chance level during training trials, but not during test trials (training: W = 45, p = 0.0039; N = 9; test: W = 13, p = 0.4492; N = 9; Wilcoxon signed rank test). The trajectories and the duration until visiting a feeder were longer for test trials than for training trials (Figures 2D, S4A; distance: W = 339, p < 0.0001; n = 27; duration: W = 325, p < 0.0001; n = 27; Wilcoxon matched pairs signed rank test). Similar results were obtained when adding a small visual pattern during test trials (Figure S1).

**Figure 2.**
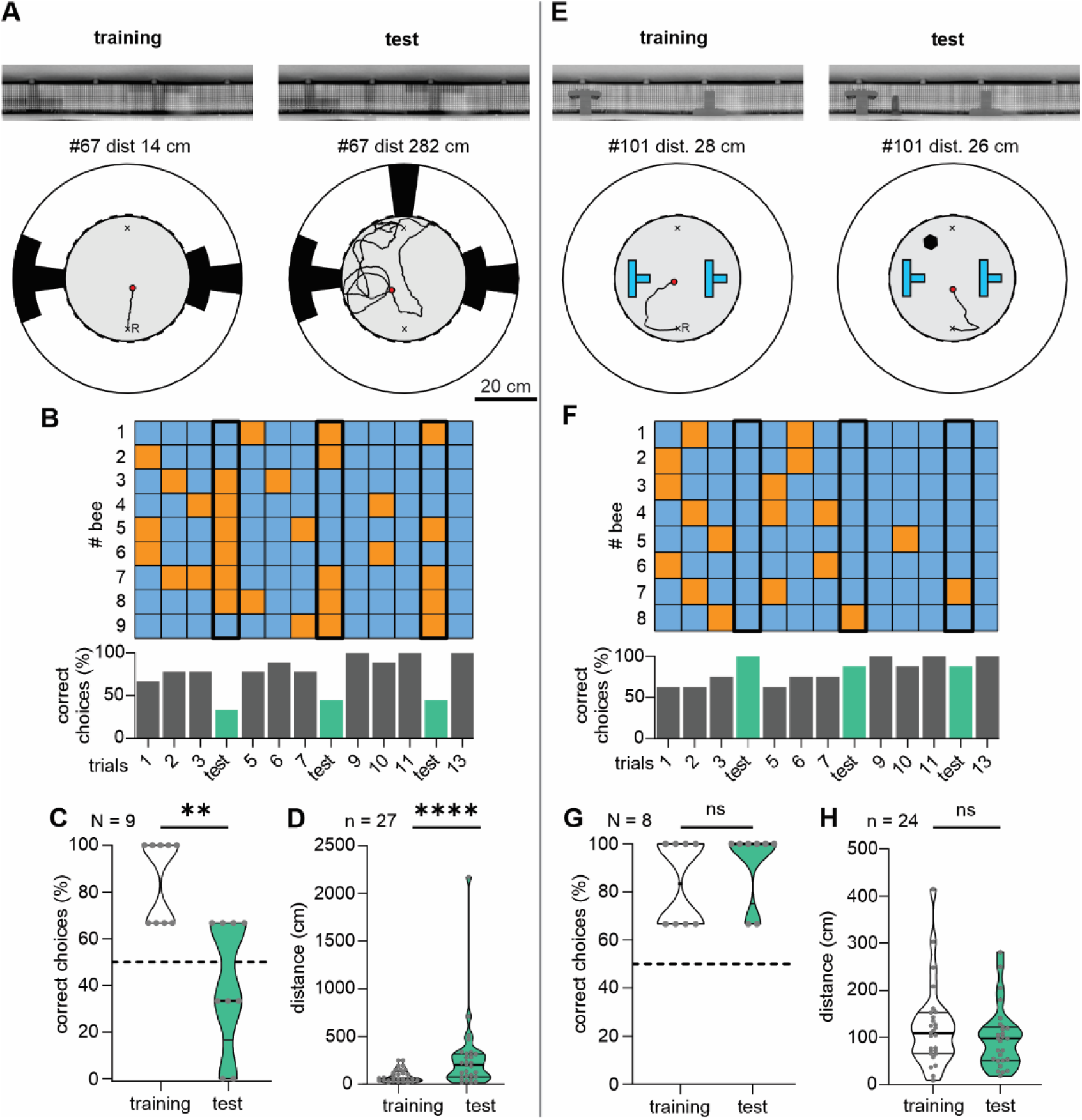
Honeybees acquire a retinotopic image for visual patterns but not when presenting local objects. (A) Panoramic photo from the arena centre and schematic top-down view of the arena during training and test. Trajectories of one example animal are presented. (B) *Top*: Colour-coded choice matrix visualizing all individual decisions (blue = correct; orange = wrong). Each row represents one honeybee. Test trials are highlighted with a thicker black frame. *Bottom*: Percentage of correct choices for all honeybees split in trials (training = grey bars; test = green bars). (C) Mean decisions for the trials and directly preceding training trials. Correct choices were higher during training than test (W = 36, p = 0,0078; N = 9; Wilcoxon matched pairs signed rank test) and were only above chance level during training (training: W = 45, p = 0.0039; N = 9; test: W = 13, p = 0.4492; N = 9; Wilcoxon signed rank test). (D) Travelled distances during test and training trials (W = 339, p < 0.0001; n = 27; Wilcoxon matched pairs signed rank test). (E-H) Same as in (A-D), but in the presence of local objects instead of visual patterns. Statistics: (G) testing against chance level for training: W = 36, p = 0.0078, N = 8 and for test: W = 36, p = 0.0078, N = 8; Wilcoxon signed rank test. Training versus test: W = 5, p = 0.6250, N = 8; Wilcoxon matched pair signed rank test. (H) Difference in distance: W = 136, p = 0.0526, n = 24; Wilcoxon matched pairs signed rank test).

### Honeybees learn three-dimensional objects differently than visual patterns

We next tested whether honeybees also learn a panoramic snapshot when presented with three-dimensional objects instead of visual patterns. We thus reconstructed the view from the previous experiment with DUPLO® bricks (Figure 2E), and a cylinder was added for test trials. The arena was homogenously illuminated by the LED-panels. For comparability, we used the same protocol as for the experiments with the visual patterns. To our surprise, in the test trials, the honeybees were not disoriented. Trajectories and choices did not differ from training trials (Figure 2E-2G; W = 5, p = 0.6250; N = 8; Wilcoxon matched pair signed rank test) and the percentage of correct choices were above chance level during both training and test trials (Figure 2G; training: W = 36, p = 0.0078; N = 8; test: W = 36, p = 0.0078; N = 8; Wilcoxon signed rank test). Consistent with these results, travelled distance and trial duration were similar between training and test trials (Figures 2H, S4A; distance: W = 136, p = 0.0526; n = 24; duration: W = 53, p = 0.4600; n = 24; Wilcoxon matched pairs signed rank test). In summary, adding a local object into a previously learned constellation of other objects did not degrade the honeybees’ ability to find the correct feeder. These findings suggest that honeybees remember objects as single landmarks, which become contrasted by the visual patterns that were stored as a coherent panoramic snapshot.

### Conflict experiment

We next asked whether honeybees rely more on a landmark-based or a panoramic snapshot memory. To this end, we set both visual cues in conflict and tested whether the animals were predominantly guided by the visual patterns or by the local landmark. For training, we presented both, the visual patterns and a local landmark (Figure 3A). We expected that the honeybees would learn the location of the correct feeder by associating the feeder with the visual patterns and with the local landmark. During training, we observed a logistic increase in correct choices over trials (Figure S3A; R^2^ = 0.6344; n = 10; N = 9). The rate of correct choices was above chance level (Figure S3B; t = 3.467, p = 0.0071; n = 10; N = 9; one sample t-test). Following three training trials, we induced a conflict between the visual patterns and the local landmark by exclusively moving the local landmark to the arena’s opposite site (*test*, Figure 3A). If the honeybees were guided by the visual patterns, then they should ignore the displaced local landmark and approach the same feeder as during training. If the honeybees, however, were guided by the local landmark, then they should approach the other feeder. Interestingly, in 23 of 27 test trials (85.2%), the bees were guided by the local landmark, a decision rate well above chance level (Figure 3B and 3C; W = 41, p = 0.0156; N = 9; Wilcoxon signed rank test). Furthermore, the honeybees showed no sign of disorientation during test trials because travelled distance and duration were similar to the preceding training trial (Figures 3D, S4B; distance: W = 66, p = 0.4375; n = 27; duration: W = 55, p = 0.5186; n = 27; Wilcoxon matched pairs signed rank test). Collectively, these results demonstrate a preference for the local landmark.

**Figure 3.**
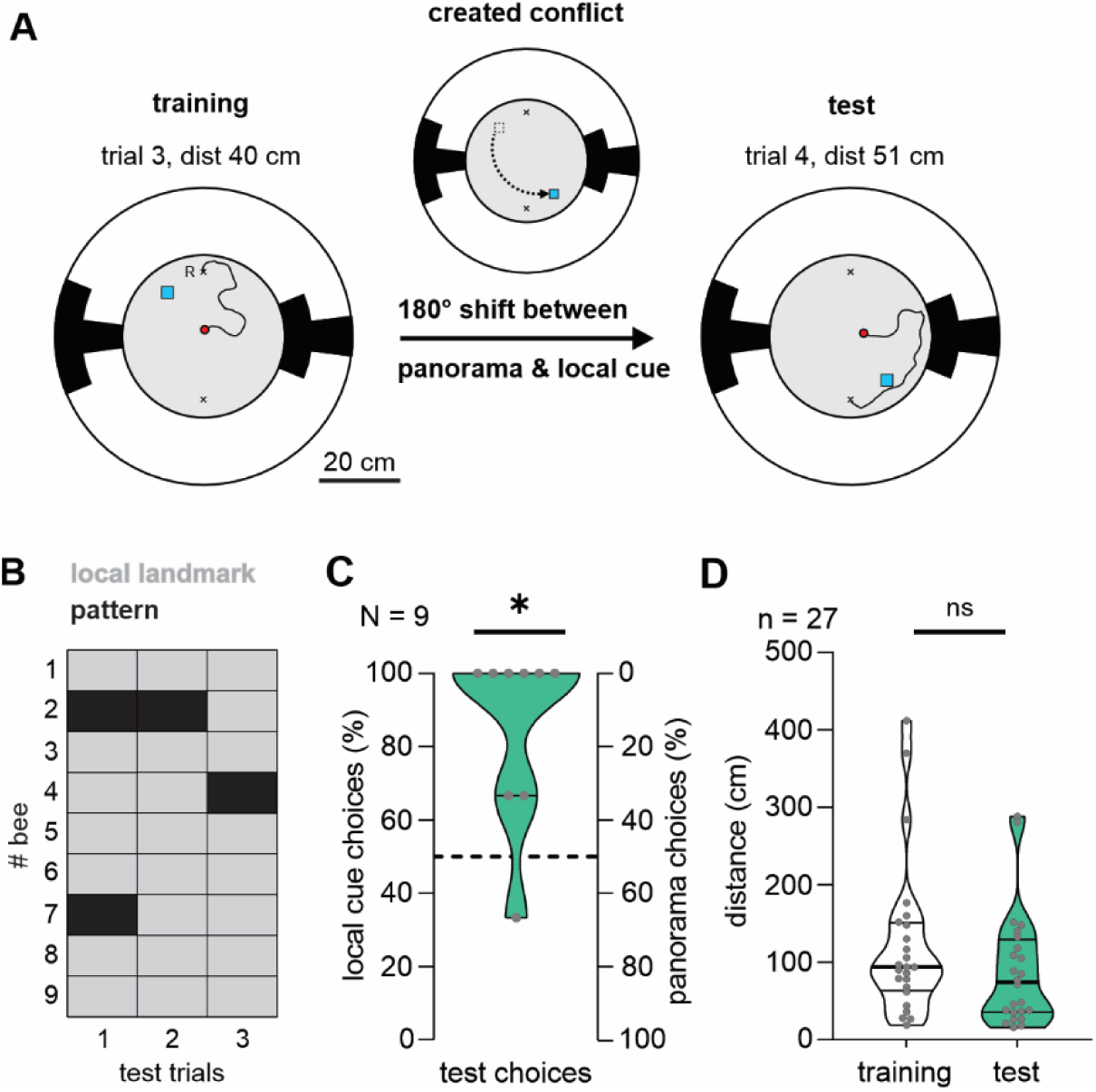
When set in conflict, honeybees rely on the local landmark for guidance. (A) Training in the presence of a panoramic pattern and a local landmark. For these tests, the local landmark and the panoramic pattern were set in conflict. (B) Individual choices for each test. Light grey is a choice for the feeder cued by the local landmark, black a choice for the feeder cued by the panoramic pattern. (C) Percentage of local landmark choices for all honeybees (W = 41, p = 0.0156; N = 9; Wilcoxon signed rank test). (D) Travelled distances for test and preceding training trials (W = 66, p = 0.4375; n = 27; Wilcoxon matched pairs signed rank test).

### Two navigational strategies: Image matching *versus* landmark-based navigation

Although our results from the conflict experiment suggest that the local landmark and the visual patterns are stored independently from each other, the results could alternatively be interpreted as the given object being part of a stored panoramic snapshot. Because of its closeness to the rewarded feeder, the object may have contributed to a substantial portion of the insect’s retinal image when the honeybee approached the rewarded feeder. Therefore, the object was weighed more than the visual patterns covering a smaller portion of the retinal image. Ultimately, to test whether the object is separately stored from a panoramic snapshot of visual patterns, we conducted a last series of experiments. In the presence of the visual patterns, naïve honeybees were either trained with two objects or with only one. Five animals were first trained with one object (*one local cue*; Figure 4A) and five additional bees with two objects (*two local cues*, Figure 4B). In training trials with two local cues, the objects were placed so that they provided ambiguous spatial information, i.e., the honeybees could not discriminate between feeders based on local cues and instead had to rely on the visual patterns. For the test trials, we added a vertical stripe to the visual pattern.

**Figure 4.**
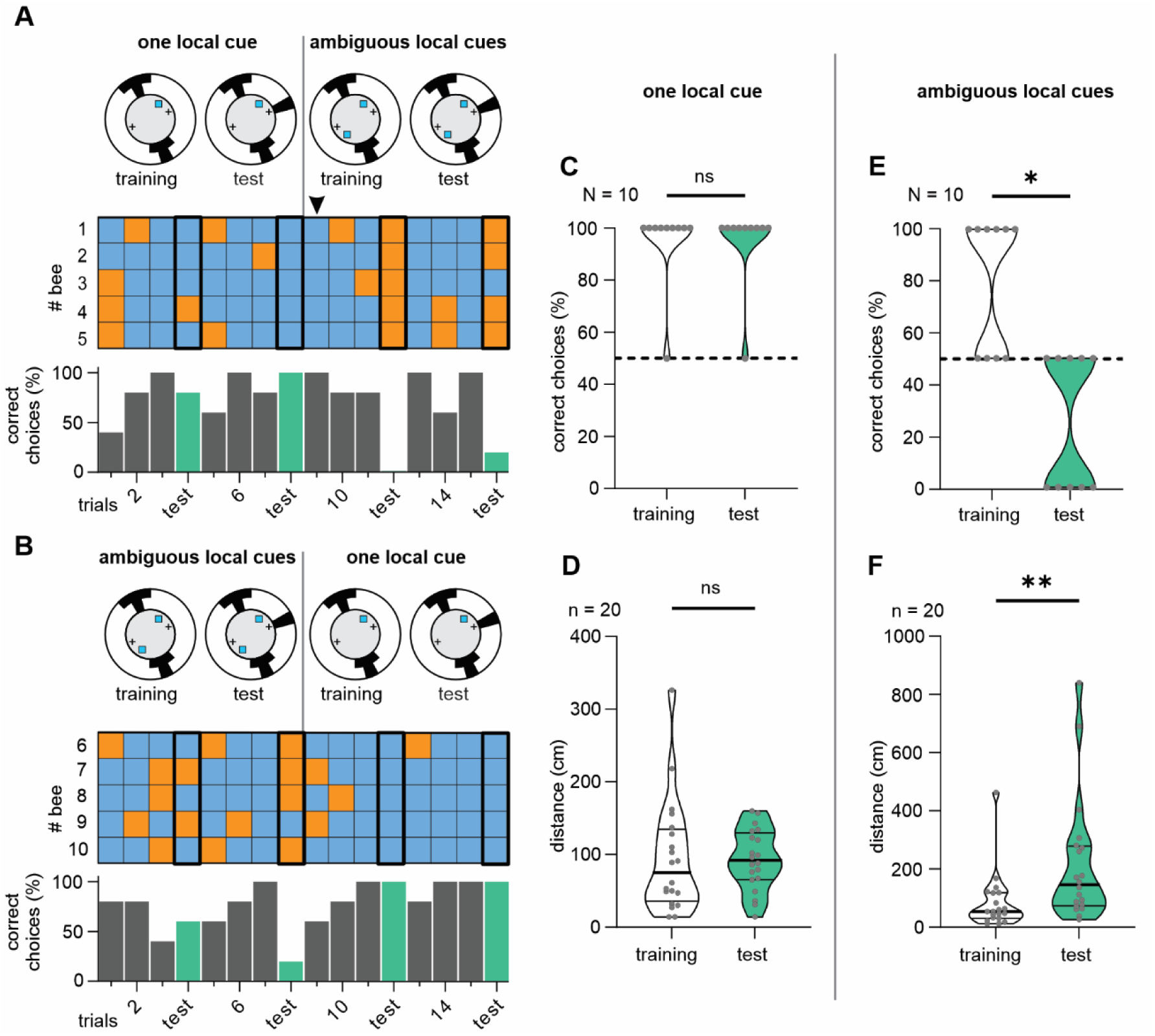
Two navigational strategies for place learning. Schematic top views of the arena during training and test, as well as the individual choices as a colour-coded matrix (blue = correct; orange = wrong). Black arrow in (A) points to the first decisions after the scenario was changed. Histograms below show the percentage of correct choices for all honeybees split in trials (training = grey bars; test = green bars). (A) One cohort of five animals started foraging either in the presence of one object (*local cue*) and the visual pattern or (B) in the presence of two ambiguously placed local cues and the visual pattern. After two tests, the training scenario was changed accordingly. (C) Percentage of correct choices when one local cue was present for training and test trials (training versus test: W = 0, p > 0.9999; N = 10; Wilcoxon matched pairs signed rank test). Percentage of correct choices differed from chance level (dashed line; training: W = 45, p = 0.0039; N = 10; test: W = 45, p = 0.0039; N = 10; Wilcoxon signed rank test). (D) Travelled distance when one local cue was present (W = 18, p = 0.7562; n = 20; Wilcoxon matched pairs signed rank test).(E) Same as (C) but when two ambiguous cues were present (training versus test: W = 28, p = 0.0156; N = 10; Wilcoxon matched pairs signed rank test; chance level: training: W = 21, p = 0.0312; N = 10; test: W = 15, p = 0.0625; N = 10; Wilcoxon signed rank test). (F) Same as (D) but when two ambiguous cues were present (W = 138, p = 0.0083; n = 20; Wilcoxon matched pairs signed rank test).

Consistent with our hypothesis, honeybees were well oriented in test trials where one local cue was present (Figure 4A, 4B). They reliably located the correct feeder (Figure 4C; training versus test: W = 0, p > 0.9999; N = 10; Wilcoxon matched pairs signed rank test; chance level: training: W =45, p = 0.0039; N = 10; test: W = 45, p = 0.0039; N = 10; Wilcoxon signed rank test) and the travelled distance and trial duration did not differ from training (Figures 4D, S4C; distance: W = 18, p = 0.7562; n = 20; Wilcoxon matched pairs signed rank test; duration: t = 0.4539, p = 0.6550; n = 20; paired t-test). Interestingly, although the honeybees could have ignored the visual patterns during the preceding training trials and learned only the spatial relationship between the object and the rewarded feeder, they chose the correct feeder from the very first training trial with ambiguously placed local objects (Figure 4A *black arrowhead*). This result demonstrates that honeybees have associated the visual patterns with the rewarded feeder in the first training block.

By ambiguously positioning two objects, we forced the honeybees to be guided by the panoramic pattern. Adding now a vertical stripe in test trials induced disorientation, represented by more visits to the wrong feeder (Figure 4E; training versus test: W = 28, p = 0.0156; N = 10; Wilcoxon matched pairs signed rank test; chance level: training: W =21, p = 0.0312; N = 10; test: W = 15, p = 0.0625; N = 10; Wilcoxon signed rank test), increased travelled distance and trial duration compared to the training (Figure 4F, S4C; distance: W = 138, p = 0.0083; n = 20; duration: W = 172, p = 0.0006; n = 20; Wilcoxon matched pairs signed rank test).

Thus, our results show that visual patterns and local landmarks are learned independently from each other. In addition, depending on which cue is more reliable, honeybees can flexibly choose which memory to rely on.

## Discussion

In this study, we designed a behavioural paradigm to test for visual spatial memory in freely walking honeybees. Consistent with the ‘snapshot’ theory of spatial memory (Judd and Collett, 1998, Zeil et al., 2003, Collett et al., 2013), adding a novel visual pattern, which exclusively perturbed the panorama, significantly impaired the honeybees’ capability to relocate the correct feeder (Figure 2C, 2D). Even adding a small visual pattern (Figure S1) evoked sufficient changes in the panorama so that no view matched well enough with previously learned views. In contrast to the visual patterns, adding an object to a set of previously learned objects did not affect the honeybees’ orientation behaviour (Figure 2G, 2H). These findings were unexpected, considering that the panoramic views were comparably affected, as in the experiments with the visual patterns (Figure S2). In essence, the change in rotational image difference was comparable between the visual scenes consisting of the patterns and the local objects. This result suggests that the different orientation behaviours, elicited by adding a novel pattern or object, cannot be explained through different impacts on the panoramic view.

Our results suggest that honeybees learn two-dimensional visual patterns differently than three-dimensional objects. Doussot et al. also showed that flying bumblebees remember local objects independently from visual patterns presented in the background (Doussot et al., 2020). In a conflict experiment, similar to ours (Figure 3), bumblebees primarily relied on visual patterns for orientation. Although the honeybees in the present study preferred a landmark-based navigation over an image-matching strategy based on visual patterns, both experiments support the idea that bees learn local objects independently from visual patterns. Furthermore, in neither case were the animals disoriented, which speaks against a single, all-encompassing snapshot memory (Wystrach et al., 2011), as the panoramic view changed greatly in response to the conflict. While the results of the conflict experiments in bumblebees could be explained with a model assuming the storage of multiple snapshots (Doussot et al., 2020), the present results, notably Figure 2E-2H, cannot be explained with a multi-snapshot strategy. Conversely, our results suggest that honeybees learn a panoramic snapshot that includes the visual patterns, while local objects are learned as single landmarks. Our last experiment further supports this interpretation (Figure 4). When honeybees were trained with both an object and a visual pattern, and the spatial information of the object was subsequently rendered ambiguous, the bees used a panoramic snapshot of the visual patterns as backup for orientation from the first trial on (arrowhead Figure 4A). This backup worked well until a novel visual pattern was introduced, which induced disorientation as predicted from the results in Figure 2A-D. When spatial information provided by the object was unambiguous, the honeybees were not disoriented in response to the novel visual pattern. Altogether, this experiment clearly shows that the honeybees had learned the object independently from the visual patterns and that they flexibly used both memories.

Which physical parameters are important to learn objects as single landmarks or as part of a panoramic image remains unclear. From the honeybees’ walking trajectories, we noticed that the bees neither had to touch nor circumvent the objects to learn them. One important aspect that differed between the objects and the visual patterns is that the objects had a three-dimensional shape and depending on the honeybee’s position and heading direction the objects in the foreground occluded different parts of the visual patterns in the background. From previous experiments, it is known that motion parallax is important for flying bees to measure the distance to objects in their surroundings (Lehrer et al., 1988, Srinivasan and Zhang, 2004) and thus to distinguish objects in the foreground from the background (Dittmar et al., 2010). Future studies are needed to address the extent to which the walking honeybees in our study could have used motion parallax to differentiate objects from the visual patterns. In line with our results, recent findings in tethered-walking bumblebees navigating a virtual reality revealed that bumblebees learn objects separately from other objects that differ in size, colour, or pattern (Eckel et al., 2025). Bumblebees were trained to remember a food location that was surrounded by two constellations of cylinders, an inner and outer one. Displacing the inner cylinders revealed that bumblebees primarily relied on these cylinders to find the reward. This finding was independent from the cylinder size and hence the proportion of the retinal image that was covered by the inner cylinders and emphasizes that insects do not exclusively learn a panoramic snapshot at the goal’s vicinity but that they are indeed able to group and remember objects independently from other objects.

Taken together, our results imply that honeybees remember two-dimensional visual patterns as a panoramic snapshot, while simultaneously learning objects in the foreground as distinct visual cues. This scenario indicates the presence of two independent memories that are processed in parallel, similar to findings from the mammalian brain (Morris, 1981, Parron et al., 2004, Renaudineau et al., 2007, Knierim and Hamilton, 2011). By monitoring the brain activity of walking honeybees that acquire spatial memory from objects and/or visual patterns, we will address these neural aspects in the future.

## Acknowledgements

We thank Patrick Schultheis, Basil el Jundi and Karen Mesce for their helpful comments on our manuscript. We thank Konrad Öchsner for building the LED arena. In addition, we thank Dirk Ahrens for beekeeping.

## Author contributions

Conceptualization: NS, KP, MJB

Methodology: NS, MJB

Formal Analysis: NS

Investigation: NS

Visualization: NS, MJB

Funding acquisition: MJB

Project administration: MJB, KP

Supervision: MJB, KP

Writing – original draft: NS, MJB

Writing – review & editing: NS, MJB, KP

## Methods

### Lead contact

Further information and requests for methods and reagents may be directed to and will be fulfilled by the lead contact, Jerome Beetz.

### Materials availability

This study did not generate new unique reagents

### Bees

We used honeybees (*Apis mellifera carnica*) that were kept by the bee-station of the University of Würzburg. At the hive entrance, we caught returning foragers, indicated by carrying pollen. Because these bees were caught at their hive, their home vector was reasoned to be zero and hence path integration should not have interfered with the animals’ performances in our experiments (Scheiner et al., 2020). Before experiments, honeybees were briefly immobilized on ice and their wings were clipped. Afterwards the animals had at least 20 min to recover. To support recovery, honeybees were given a small amount of 60% sucrose solution. Each honeybee was used for only one experiment.

### Experimental setup

We used a cylindrical arena with a diameter of 40 cm and a height of 20 cm that was placed inside the laboratory. The inner wall of the arena was covered with evenly spaced 16 × 128 RGB-LEDs (M160256CA3SA1, iPixel LED Light Co., Ltd, Baoan Shenzhen, China). The LED-screen covered 360° horizontally and 45° vertically, when viewed from the centre with a horizontal angular resolution of 2.8°.

The LEDs were controlled with a Raspberry Pi (Model 3B, Raspberry Pi Foundation, Cambridge, UK). Light intensity was measured with a spectrometer (Maya2000 Pro, Ocean Optics) for each quadrant from the centre of the arena at the typical eye level of a honeybee and averaged. For our ‘T’-shaped pattern the intensity was at 1.09 × 10^15^ photons/cm^2^/s (Figure 1), with the addition of the vertical stripe it was 1.08 × 10^15^ photons/cm^2^/s (Figure 2A – D, *test*), with the addition of a smaller spot it was 1.14 × 10^15^ photons/cm^2^/s (Figure S1), when using local Landmarks instead of the pattern it was 9.97 × 10^14^ photons/cm^2^/s (Figure 2E – H, *training*), when adding a novel landmark to the local landmarks it was 1.03 × 10^15^ photons/cm^2^/s (Figure 2E – H, *test*), for the conflict experiment between pattern and landmark it was 1.14 × 10^15^ photons/cm^2^/s (Figure 3) and when presenting both pattern and two local landmarks the intensity was at 1.09 × 10^15^ photons/cm^2^/s (Figure 4).

The floor of the arena was a white-painted aluminium sheet (Firstlaser GmbH, Bardowick Hamburg, Germany) with a grid of evenly spaced holes, 4.5 cm apart from another, and had a diameter of 0.5 cm.

To occlude any visual cues outside the arena, the arena’s top was covered with a wooden lid. The lid had a camera (ELP 1080P, Shenzhen Ailipu Technology Co, China) installed in the centre to record all trials at a resolution of 1920 × 1080 px at 30 fps.

To study the honeybees’ orientation behaviour, we designed a two-choice paradigm where the animals had to choose between two identical feeders that were placed opposite from each other. The feeders consisted of a small tube that was guided through one of the holes in the arena’s floor. Attached to the tube was a small syringe that was either filled with sucrose solution (60%) or distilled water. With this setup, we could manually adjust the level of the solution from outside the arena without interfering with the experiments. This setting allowed us to manually lower the level of the sucrose solution, as soon as the honeybees had druck from it for a few seconds. Inside the arena we made sure that the tubes of both feeders only stuck out a few millimetres. We placed a small piece of orange coloured paper with a hole in the centre on top of the feeders to cue them. We always made sure that the two feeders looked identical. The feeders were 16 cm away from the centre of the arena. During training trials, only one feeder was rewarded with a 60 % sucrose solution, during tests both feeders were filled with distilled water.

### Visual stimuli

Visual patterns were created with the LED panels covering the inner walls of the arena. To present an upright ‘T’ and an upside-down C in black, 480 LEDs (240 for each T-shape) were tuned off while the remaining LEDs were white illuminated. Both ‘T-shapes’ were set orthogonally to the feeders. The ‘novel shape’, which we added in test trials in the second and fourth experiment, was a vertical stripe placed orthogonally to the ‘T-shapes’ (Figure 2A). This stripe was presented by turning 96 LEDs off. The smaller spot used for the experiment in Figure S1 was presented at mid-level elevation and covered 24 LEDs.

For some experiments, we placed real, three-dimensional objects as landmarks (‘local landmarks’) inside the arena. For the second experiment we used blue DUPLO® bricks to recreate the ‘T’ shapes from the panoramic pattern (not to scale) which had heights of 8 cm (Figure 2E). Like the panoramic pattern, these landmarks were placed orthogonally to the feeders. They were 12 cm away from the centre, which gave enough space to be circumvented by the honeybees. For test trials, we added a black metal cylinder with a height of 5.5 cm. The cylinder was randomly presented at 8 cm next to either the rewarded or the unrewarded feeder. When presenting the local landmarks alone, the LED arena was homogenously illuminated by setting all LEDs to white.

For experiment 3 (Figure 3), we used a blue straw, stabilized by a metal rod, with a small white marble at the top. This landmark had a height of 11.5 cm and was placed either to the left or the right of the rewarded feeder, 13 cm away.

For experiment 4 (Figure 4), we used either one or two identical landmarks made of blue DUPLO® bricks, with a height of 6.5 cm and a distance to the feeder of 8 cm.

## Experimental protocol

### Pre-training

To give the honeybee a chance to familiarize herself with the arena, we started with a short pre-training, where the animal was allowed to walk freely in the arena. During this time, the visual cues that were used for the subsequent training were already present. Ideally, during this time, the honeybee found the rewarded feeder by herself, which marked the end of the pre-training. If she did not locate the rewarded feeder within five minutes, we gently guided her towards it by using a small brush. Once the animal was at the rewarded feeder, we let her drink for 3-5 seconds before catching her with an opaque plastic cup and placing her back to the centre of the arena. The honeybee was then kept under the cup till the first trial was started. For follow-up experiments we only used honeybees that drank from the rewarded feeder.

### Experiment 1

This experiment tested whether honeybees could learn the location of the rewarded feeder by using the ‘T’-shape pattern on the LED screen for orientation. To this end, an individual honeybee was released at the centre of the arena by lifting an opaque cup. Afterwards, the honeybee walked freely in the arena until she decided for one of the feeders. We defined a choice only when the bee was probing i.e. extended its proboscis. If she found the rewarded feeder, we let her drink for 3-5 seconds. This period was short enough to avoid losing her motivation to forage. If the honeybee tried drinking from the non-rewarded feeder, she was recaptured and placed back to the arena’s centre. To ensure that the bee only used our presented visual stimuli for navigation, we randomly rotated the visual cues by 90° clockwise or counterclockwise or by 180° in conjunction with the feeders, when the bee was under the opaque cup. The floor in contrast was not rotated. In essence, only the visual cues were consistently spatially related with the feeders. Other cues, such as visual cues outside the arena or any olfactory cues, such as scent marks the honeybee could have deposited in previous trials, would not have been reliable. Afterwards, the animal was released for the subsequent trial. To be included for the analysis, each honeybee had to perform at least twelve trials.

### Experiment 2

In this experiment we implemented test trials in addition to the training trials. These tests were performed after three training trials each. In total, each animal had to perform three tests and ten training trials. During tests no reward was present. During training always one feeder was rewarded and its spatial relation to the presented stimuli was consistent.

For the first part of experiment 2 we presented the ‘T’-shape panoramic pattern during training. During tests only, we added the novel shape into the pattern (Figure 2A). This way we could analyse its effect and compare it to the training condition within the same animal. The novel shape was randomly either behind the previously rewarded or unrewarded feeder.

For the second part we used the local ‘T’ landmarks during training and implemented the novel landmark in tests just like in the first part. We used different animals for each part of the experiment.

### Experiment 3

In the third experiment we presented the ‘T’ shape panoramic pattern as well as a blue straw as an additional local landmark that could be used for guidance. We used the same trial order as in experiment 2. During tests the spatial relation between the panoramic pattern and the local landmark was shifted by 180° creating a conflict as the panorama now indicates one feeder, while the landmark indicates the other. Thus, for this experiment specifically there was no correct and incorrect choice during the tests, but rather a choice for either the panorama or the local landmark for guidance.

### Experiment 4

For the last experiment we had two different conditions for each animal. In one case we presented the panoramic pattern as well as a local landmark. During the tests we additionally showed the novel pattern we used in experiment 2. In the other case we once again presented the panoramic pattern but now together with two identical landmarks, one at each feeder, making them ambiguous and thus useless for guidance. During tests, we would again show the novel pattern in the panorama. In both cases, honeybees had to perform six training trials and two tests. Importantly, every animal was presented with both cases, half of the animals starting with one local landmark, the other half with the two ambiguous landmarks (Figure 4A and 4B).

### Analysis

We noted the individual choices of each honeybee, which was confirmed offline based on video recordings. To analyse the animals’ orientation behaviour, we computed walking trajectories from the recorded videos. For tracking, we used the software Ctrax (version 0.5.18, Caltech ethomics project, (Branson et al., 2009)) which outputs xy-coordinates for each individual frame. Tracking errors were corrected with the associated MATLAB scripts (data_analysis_trajectories.m; downloaded and modified from: ctrax.sourceforge.net).

For experiment 1, we quantified the percentage of correct choices, i.e., choices for the rewarded feeder across trials for ten honeybees.

For the remaining experiments, we compared the performance between the test trials and the directly preceding training trial training. We compared the differences in the mean percentage of correct choices, travelled distances and the trial duration. For experiment 3, we analysed the test choices regarding the percentage of local cue choices i.e. how often honeybees chose the feeder indicated by the local landmark.

To quantify the visual scene, we took 360° pictures (5376 × 2688 px) from the centre of the arena for each experimental condition (camera: RICOH THETA SC2, Tokyo, Japan). Using a custom MATLAB script (image_correlation.m) we then grey-scaled and down-sampled the images via a bilinear interpolation to 1 px/deg (180 × 359 px) to match with the honeybee’s estimated visual resolution (Stürzl and Zeil, 2007). To see how adding a novel landmark in test trials affects the visual scene from the training trial, we computed the root mean squared error between the images that were taken before and after adding the novel landmark (Figure S2A; (Narendra et al., 2013, Narendra and Ramirez-Esquivel, 2017)). In addition, we compared the image differences between the original image and the ones created by rotating the original image in steps of 1° (rotational image difference function; Figure S2B). This let us estimate how an orientation based on view-matching could be realized when the honeybee rotates around her horizontal body-axis.

### Statistics

For the statistics and creating the figures, we used MATLAB (version R2022b, MathWorks, Natick, MA, USA) and GraphPad Prism (version 9.0.0 for Windows, GraphPad Software, San Diego, California, USA).

To determine a learning effect over trials, we used a logistic regression model (dose-response model with variable slope) which represents a standard model to determine learning in a two-choice paradigm (Hartz et al., 2001, Bewick et al., 2005, Pretot et al., 2016, Viering and Loog, 2023).

The equation is based on the four-parameter logistic (4PL) model. Logistic Model (4PL) Formula:

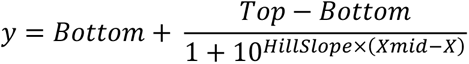

- **y** is the predicted percentage of correct choices (the dependent variable).
- **X** is the **trial number**, the independent variable (input).
- **Bottom** is the **lower asymptote**, the baseline performance (50% for random choice).
- **Top** is the **upper asymptote**, the maximum achievable performance (100% for a correct choice).
- **HillSlope** is the **slope factor** (steepness of the curve), determining how quickly the learning improves.
- **Xmid (EC50)** is the **point of inflection** where the model predicts 50% of the maximum performance, corresponding to the trial number at which the learning rate is greatest.

**R^2^** (coefficient of determination) indicates how well this model fits to our data, ranging from 0.0 to 1.0. A value of 1.0 indicates that the regression curve perfectly predicts the data points.

For the statistics, we first tested whether our data were normally distributed according to the Anderson-Darling test.

For analysing mean performance of a population, we compared whether it differed from chance level (50%) using either one sample t-tests, in case of a normal distribution, or Wilcoxon signed rank tests, if the data were non-normally distributed. These tests were always two-tailed.

When comparing data sets, as we did for comparing training and test performances and walking distances, we used paired t-tests in case of a normal distribution, or Wilcoxon matched pairs signed rank tests, if not. These tests were always two-tailed.

Significance levels are always indicated via asterisks: each additional asterisk indicates an order of magnitude increase in significance (p < 0.05 *, p < 0.01 **, p < 0.001 ***, p < 0.0001 ****).

## Supplemental information

**Video S1. Typical trial of a trained honeybee**

A typical trial of a honeybee trained to use the two-dimensional pattern (Figure 1). During this training the rewarded feeder was rapidly located by the trained animal.

**Video S2. Comparison of early and later trial**

First trial of a honeybee presented with the pattern (Figure 1) and a later trial of the same animal.

**Figure S1.**
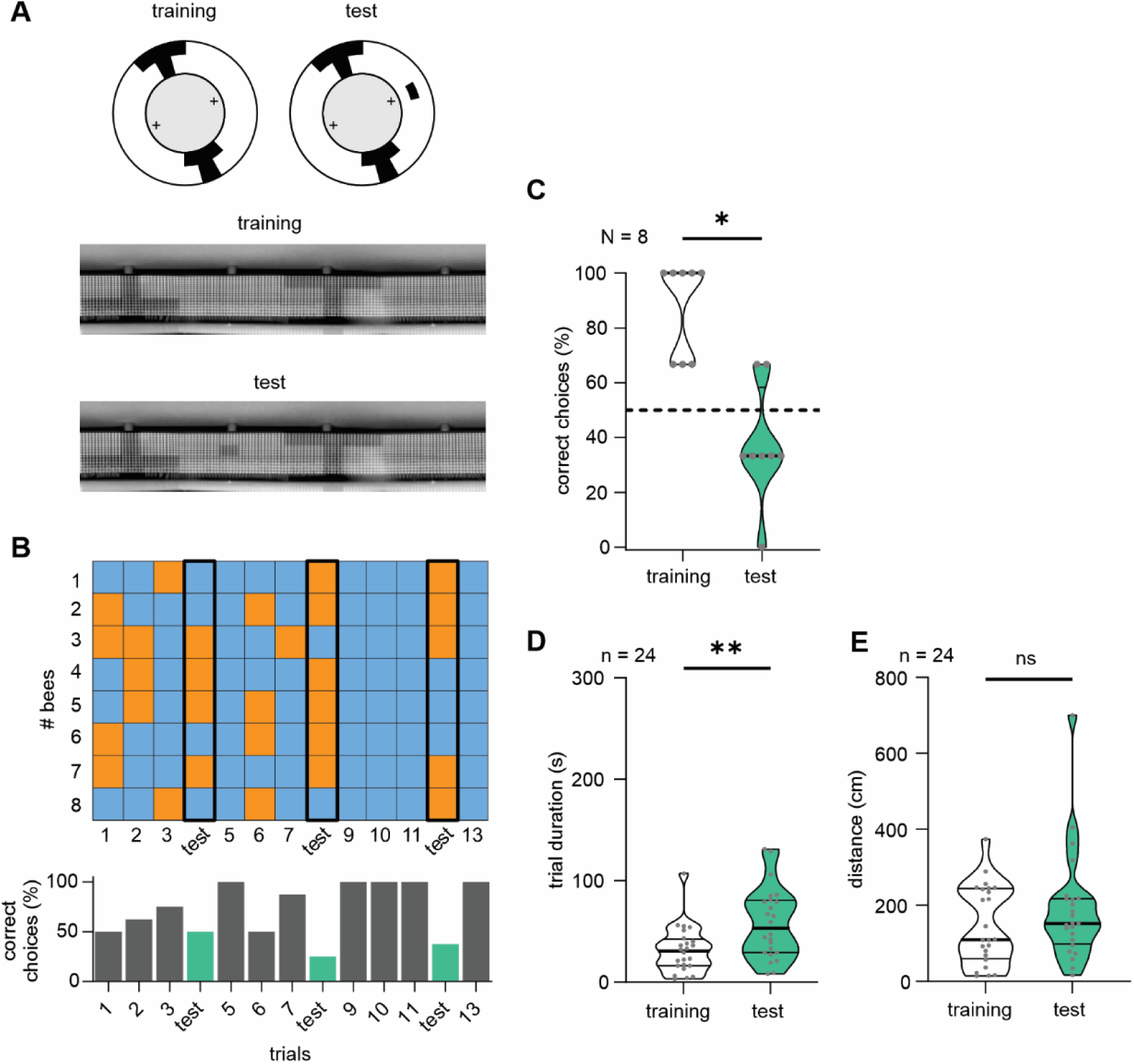
Retrieving spatial memory is affected when perturbing the panoramic view with a small visual pattern. (A) Schematic top views and down-sampled panoramic images showing the visual scene during training and test. (B) *Top*: Colour-coded choice matrix visualizing all individual decisions (blue = correct; orange = wrong). Each row represents one honeybee. Test trials are highlighted with a black frame. *Bottom*: Percentage of correct choices for all honeybees split in trials (training = grey bars; test = green bars). (C) Comparison of mean decisions for all tests, compared to their preceding training trials. Only correct training decisions were above chance level (*training*: W = 36, p = 0.0078; N = 8; *test*: W = 20, p = 0.2344; N = 8; Wilcoxon signed rank test). Rate of correct choices during training and test differed significantly (W = 28, p = 0,0156; N = 8; Wilcoxon matched pairs signed rank test). (D) Trial duration during training and test trials (t(23) = 3.392, p = 0.0025; n = 24; paired t test). (E) Travelled distances during training and test trials (W = 46, p = 0.5224; n = 24; Wilcoxon matched pairs signed rank test).

**Figure S2.**
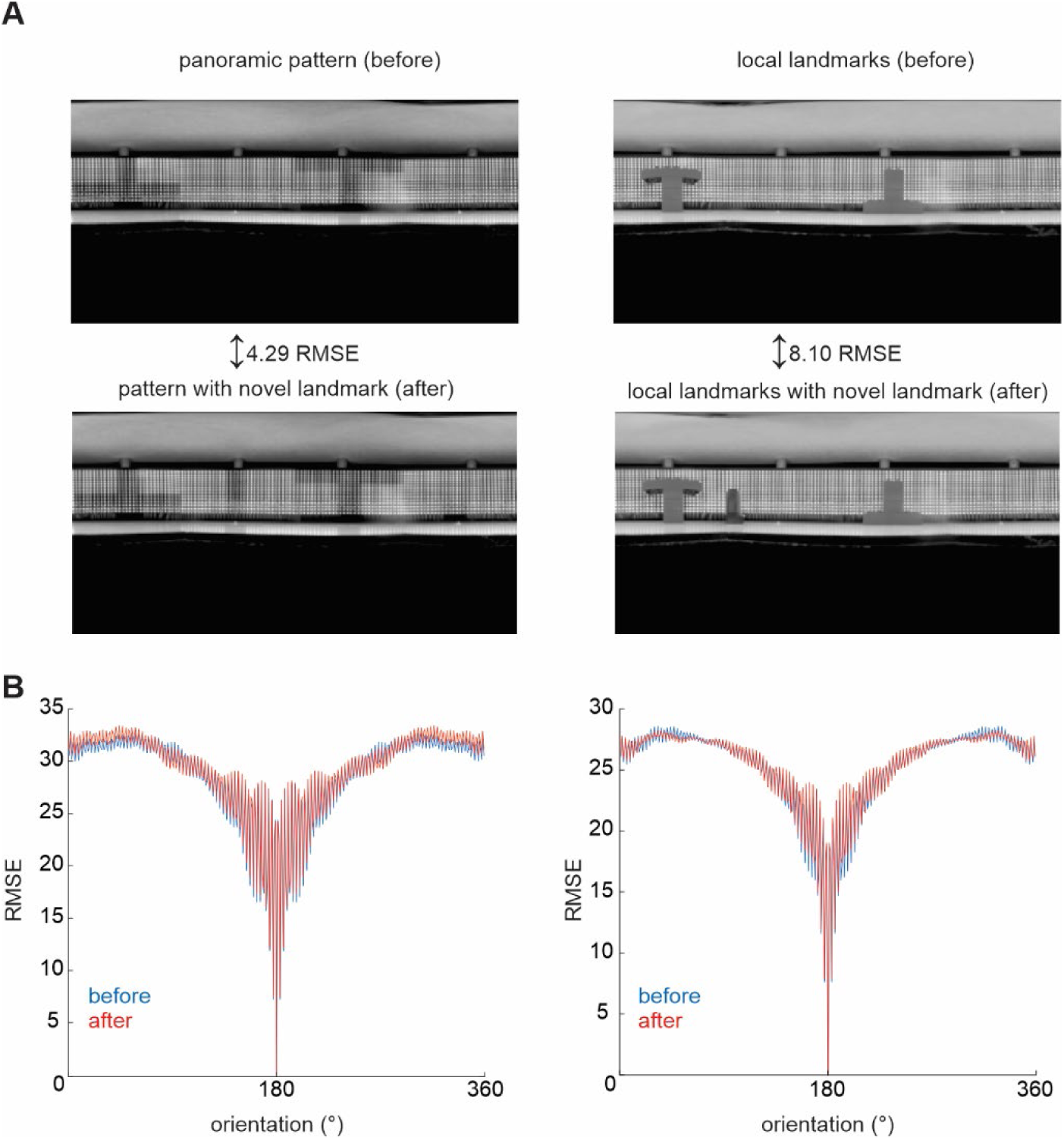
Panoramic visual scenes. (A) Panoramic, down-sampled images taken before (training) and after adding a novel landmark (test). Images were taken from the arena’s centre, the approximate release site. Root mean squared error (RMSE) quantifies the image differences between training and test situation. RMSE can range from 0 to 255, where lower values mean higher image similarity. (B) Rotational image differences were computed in steps of 1°, for the visual scenes before (*blue*) and after (*red*) adding a novel landmark. Notably, adding a landmark did not substantially change the rotational image differences neither in the case for visual patterns (*left*) nor for local landmarks (*right*).

**Figure S3.**
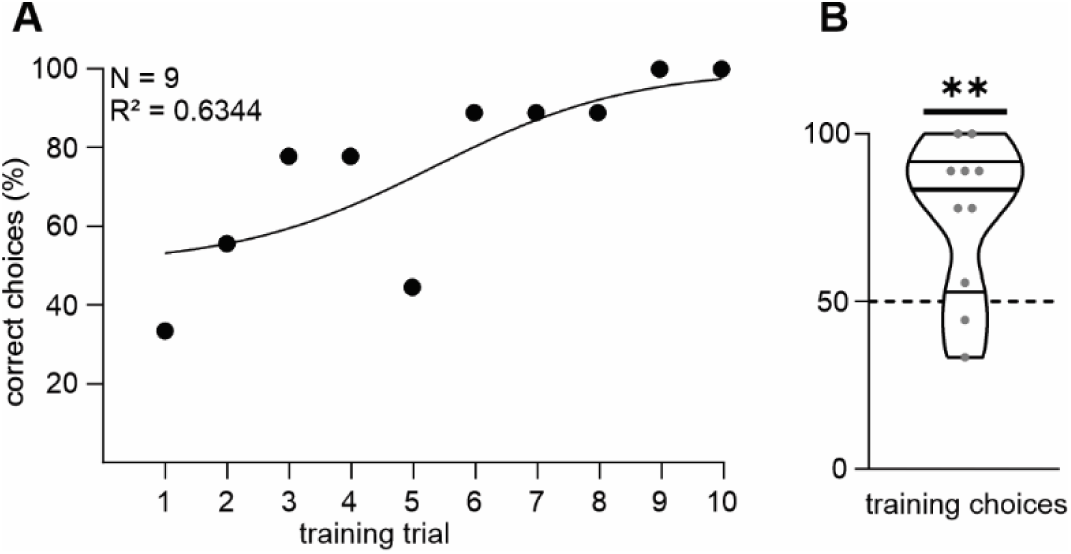
Honeybees show visual place learning when both panoramic pattern and a local landmark were present. (A) Mean training performance of ten honeybees fitted to a logistic model (N = 9; R^2^ = 0.6344). (B) Mean training choices. Dashed line indicates chance level (p = 0.0071, N = 9; one sample t-test).

**Figure S4.**
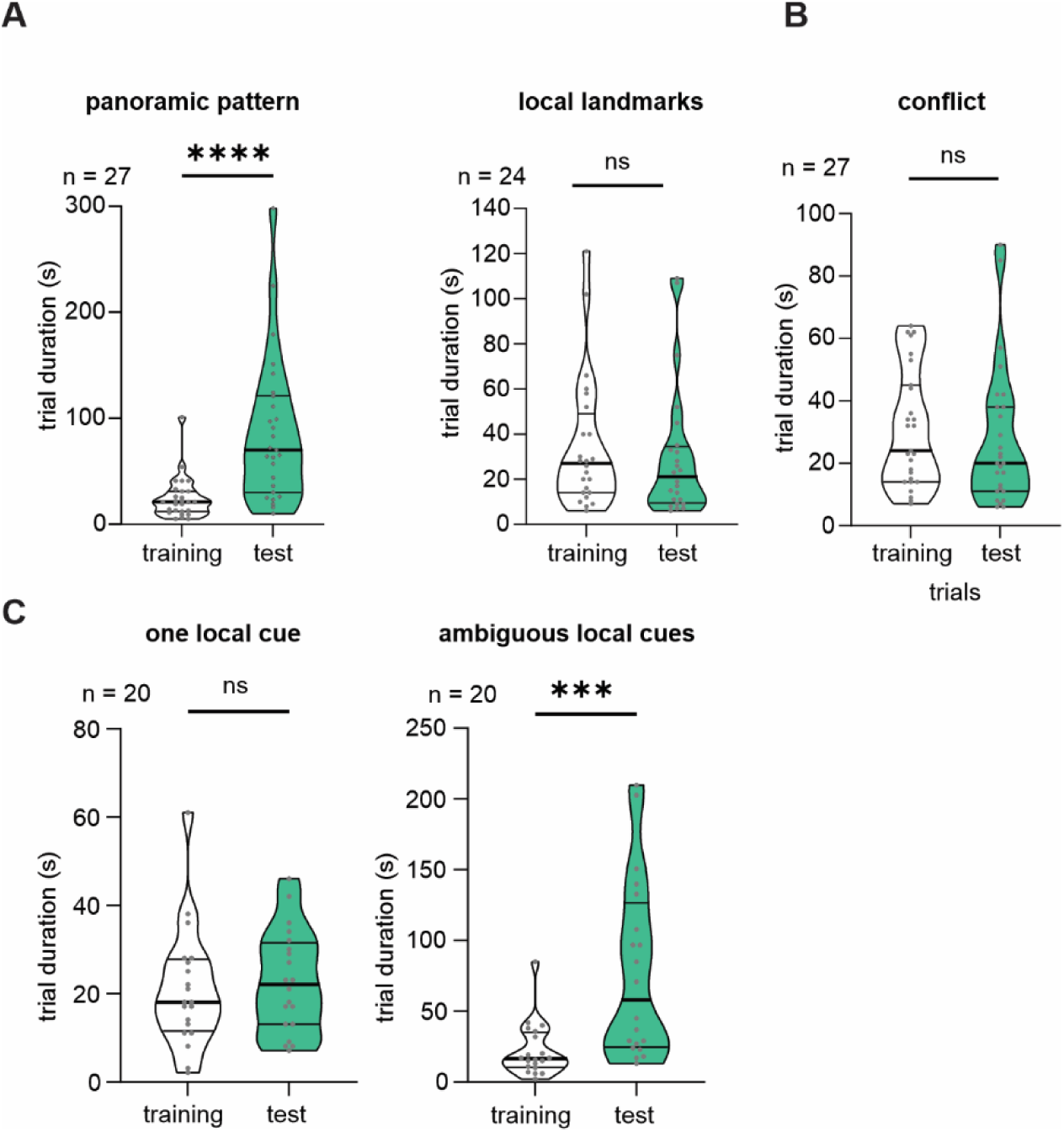
Comparison of trial durations during training and test from all experiments. Trial duration comparison between tests with their preceding training trials for experiment 2 ((*A*); pattern: W = 325, p < 0.0001; n = 27; local landmarks: W = 53, p = 0.4600; n = 24; Wilcoxon matched pairs signed rank test), experiment 3 ((*B*); W = 55, p = 0.5186; n = 27; Wilcoxon matched pairs signed rank test) and experiment 4 ((C); one landmark: t (19) = 0.4539, p = 0.6550; n = 20; paired t test; ambiguous landmarks: W = 172, p = 0.0006; n = 20; Wilcoxon matched pairs signed rank test).

